# A microfluidic platform for extraction and analysis of bacterial genomic DNA

**DOI:** 10.1101/2024.10.17.618837

**Authors:** Alex Joesaar, Martin Holub, Leander Lutze, Marco Emanuele, Jacob Kerssemakers, Martin Pabst, Cees Dekker

## Abstract

Bacterial cells organize their genomes into a compact hierarchical structure called the nucleoid. Studying the nucleoid in cells faces challenges because of the cellular complexity while *in vitro* assays have difficulty in handling the fragile megabase-scale DNA biopolymers that make up bacterial genomes. Here, we introduce a method that overcomes these limitations as we develop and use a microfluidic device for the sequential extraction, purification, and analysis of bacterial nucleoids in individual microchambers. Our approach avoids any transfer or pipetting of the fragile megabase-size genomes and thereby prevents their fragmentation. We show how the microfluidic system can be used to extract and analyze single chromosomes from *B. subtilis* cells. Upon on-chip lysis, the bacterial genome expands in size and DNA-binding proteins are flushed away. Subsequently, exogeneous proteins can be added to the trapped DNA via diffusion. We envision that integrated microfluidic platforms will become an essential tool for the bottom-up assembly of complex biomolecular systems such as artificial chromosomes.

## Introduction

The 3D spatial structure of genomes is important for gene expression and other cellular functions.^1^ Whereas eukaryotes organize their genomic DNA in a cell nucleus where individual chromosomes occupy territories^2^, bacteria organize their DNA into a compact structure called the nucleoid,^3,4,5^ which is not enclosed by a nuclear membrane. Despite much research, we still have an incomplete understanding of the 3D organization of the bacterial genome and its effects on various biological processes.^6^ There are many fruitful techniques for studying genome organization in cells such as chromosome conformation capture (3C/HiC),^7,8^ high-resolution fluorescence microscopy,^9,10^ and fluorescence-based localization techniques like FISH.^11^ Yet, many questions remain due to the inherent complexity of the cellular environment. *In vitro* single-molecule techniques are powerful since they can study DNA-proteins at the single molecule level in controlled environments, but they typically use short DNA molecules that are orders of magnitude smaller than bacterial genomes.^12,13,14^ Recently, we have proposed a novel *in vitro* method (“genome-in-a-box”) to study chromosome organization from the bottom up using purified bacterial chromosomes^15^, i.e. using DNA molecules of similar size to the genomes of living cells. Extraction of nucleoids from bacteria is nontrivial, although first examples of nucleoid isolation from bacteria date from the 1970’s.^16^ While we recently presented a method to obtain deproteinated DNA of megabasepair length from *E. coli*,^17^ it remains challenging to avoid unwanted DNA damage that occurs due to mechanical shearing during pipetting. A microfluidic system could provide solutions to these limitations, as a precise and well-defined control of fluid flow minimizes the shear forces on the megabase-scale DNA. Furthermore, confining the DNA in microscale compartments allows for continuous monitoring of individual DNA objects. Microfluidic devices have been extensively used for trapping live cells^18^ and cell-like synthetic compartments^19,20^ and so-called ‘mother-machine’^21^ devices were developed for studying the growth and controlled cell lysis^22^ of bacterial cells.

In this paper, we introduce a microfluidic platform that enables all the individual steps needed for lysis of individual bacterial cells, extraction of the bacterial nucleoid, deproteination of the nucleoid, imaging analysis of the extracted nucleoids, and introduction of DNA-structuring elements to the genomic DNA (Fig. 1). Notably, this approach allows for continuous tracking of the individual nucleoids in discrete microchambers that are hydrodynamically isolated from a buffer channel, which eliminates shear forces on the fragile genomic DNA molecules while allowing for addition and exchange of DNA-binding proteins. Flow control is provided by pneumatically actuated on-chip valves.^23^

**Fig. 1.**
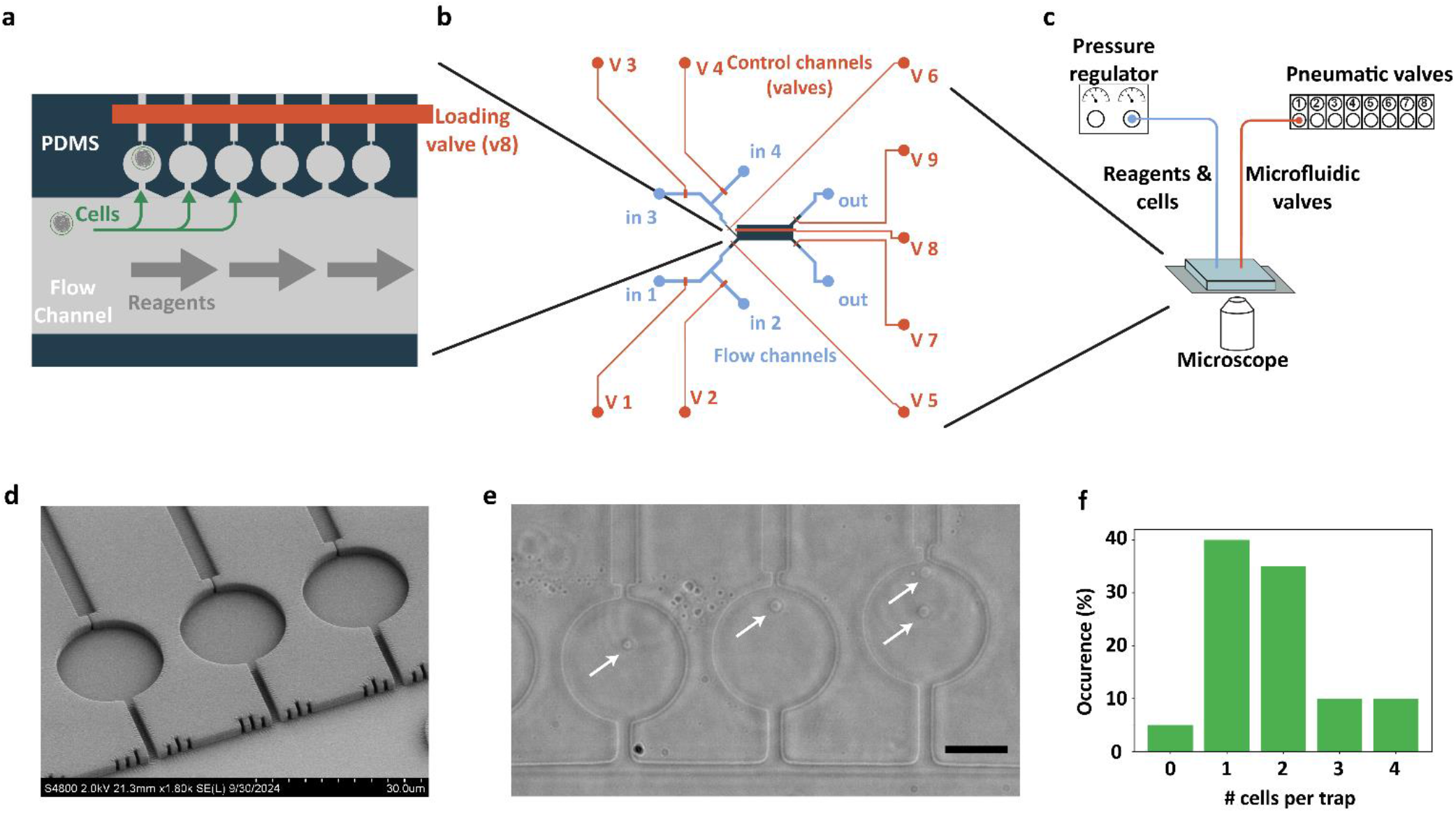
A microfluidic platform for extraction and purification of bacterial nucleoids. **a**. A liner array of microfluidic trapping chambers. Cells are inserted into the chambers by directing fluid flow from the large filling channel through the trapping chambers and out of the respective exhaust channels. The exhaust channels are too narrow for cells to pass through, allowing them to stay trapped in the chambers. **b**. Overview of the design of the microfluidic chip. Pneumatically actuated Quake valves are used in a push-down configuration to direct the flow of cells and reagents. **c**. Overview of the setup for 2-layer microfluidic platform with on-chip flow control. **d**. SEM micrograph of the PDMS trap array. **e**. Widefield micrograph of *B. subtilis* spheroplasts (arrows) in the trapping array. Scale bar is 10 μm. **f**. Average number of spheroplasts per trap. In a typical experiment, almost 40% of the traps (n=20) contain a single spheroplast.

We validate our microfluidic platform with the extraction and analysis of bacterial chromosomes of *B. subtilis* cells. Using confocal fluorescence microscopy, we can track individual cells from the moment they are inserted into the chambers, whereupon we observe their lysis, followed by deproteination, expansion, and relaxation of their chromosomal DNA. As proof-of-principle experiments of first steps towards the bottom-up assembly of an artificial chromosome, we show the effect of DNA-binding protein Fis and PEG on the 3D structure of isolated megabasepair-long DNA.

## Results

### Design of a microfluidic platform for bacterial DNA extraction

The main objective of our microfluidic platform is to perform bacterial nucleoid extraction and long-term analysis in individual micro-chambers with minimal perturbation of the chromosomal DNA. The microfluidic device is required to switch between a number of different input solutions/fluids to perform the individual steps of bacterial DNA extraction, analysis and reagent addition, while keeping the megabase-scale DNA fixed in the trapping chambers. Our initial tests revealed that megabase-scale DNA molecules are highly sensitive to the shear forces caused by flow rate fluctuations in microfluidic channels. Typically, the volume of fluid within the feeding tubes connected to a microfluidic chip is orders of magnitude larger than the volume of the microfluidic chip itself. Therefore, fluctuations in the flexible tubing led to very substantial fluctuations in the flow rate within the microfluidic chip. With these considerations in mind, we reasoned that to be able to reliably switch between different input reagents without perturbing the megabase-scale DNA molecules, all flow control would have to be incorporated into the microfluidic chip. Therefore, we chose to use PDMS/glass for the material of the microfluidic chip as it enables straightforward implementation of on-chip flow control using pneumatically actuated microvalves (Supplementary Fig. 2).^23^ This approach eliminates the dead volume effects of the connectors and tubing because the fluid flow is manipulated via integrated valves instead of external valves or syringe pumps.

Initially, we designed more conventional microfluidic trapping devices with a 2D grid arrangement of microfluidic traps (Supplementary Fig. 1). This configuration worked well for cell trapping and for their lysis (Supplementary Video 1), but keeping the extracted nucleoids localized in the traps proved to be impossible during reagent addition, since the flexible DNA polymer would inevitably exit the traps due to the applied flow (Supplementary Video 2). Therefore, we switched to a linear array of micro-chambers with individual input and output channels (Fig. 1a). The input channels of all these chambers are connected to a single ‘filling channel’ that runs parallel to the trapping array, while the output channels are actuated with a single pneumatic on-chip valve. The advantage of this ‘side chamber’ configuration is that it allows for reagents to be added to the chambers using two methods, either via direct flow or via diffusion from the filling channel. While the latter, importantly, avoided any shear forces on the fragile genomic DNA molecules while allowing for addition and exchange of DNA-binding proteins, the former, flow-based filling, was mainly used to insert the bacterial cells into the chambers. The dimensions of the input and output channels were selected such that cells could freely flow through the input channels with a width of 2 μm but would not pass through the output channel with a width of only 0.7 μm, resulting in their entrapment in the cylindrical chambers.

The input channels that connect the filling channel to the trapping microchambers were chosen as wide and short as possible to allow efficient diffusion while still providing enough insulation in order to prevent fluid flow from reaching the chamber and perturbing the DNA. The trapping chambers were 1.6 μm in height and 16 to 20 μm in diameter. The input and output channels that run through the pneumatically actuated valves had a rounded profile and a height of 10 μm (Fig. 1b, c). A detailed description of the valve design is given in Supplementary Fig. 2.

### Microfluidic side chambers enable the isolation and study of megabasepair DNA without shear flow

The process of nucleoid extraction and analysis on our microfluidic platform consists of the following steps: 1) preparation of bacterial spheroplasts; 2) injection and trapping the spheroplasts in microchambers; 3) lysis of the spheroplasts which yielded to extraction of the DNA and the disassembly of DNA-binding proteins; and possibly 4) the addition of reagents of interest for follow-up biophysical studies.

Spheroplasts are spherical-shaped bacteria of which the outer cell wall has been removed. Preparation of the spheroplasts was performed in a cell-culture flask using lysozyme to digest the bacterial cell wall. The main reason for preparing the spheroplasts outside the microfluidic device is that the spherical shape and lack of motility makes the spheroplasts much easier to trap compared to the intact cells, which can swim out of the traps. Furthermore, this approach avoids contaminating the trapping chambers with lysozyme and cell-wall degradation products. Spheroplasts were injected into the filling channel of the microfluidic device (Materials and Methods, Fig. 1a). The exhaust channels were then opened, directing the flow through the microfluidic side chambers such that spheroplasts were trapped in them (Fig. 1e).

We characterized the trapping efficiency of the system using *B. subtilis* spheroplasts. In a typical experiment, approximately 40% of the traps contained a single spheroplast and were therefore suitable for further analysis (Fig. 1e, f, Supplementary Fig 3). In the current configuration, often more than one cell was observed to enter a chamber. The efficiency can potentially be improved by optimization of the geometry of the narrow output channels such that a single cell would block the flow and thus prevent successive cells from entering the same chamber. When the desired amount of spheroplasts was inserted into the trapping chambers, the flow through the traps was stopped and the cells were ready for lysis.

We explored two methods for cell lysis, (i) based on surfactants and (ii) based on osmotic shock. For (i), we used a lysis buffer solution (Materials and Methods) containing 5% surfactant (IGEPAL) and 500 nM of the intercalating fluorescent dye (Sytox Orange) that stains DNA, to detect the chromosomal DNA. When lysis buffer was flowed into the filling channel of the microfluidic device, the trapped *B. subtilis* spheroplasts abruptly ruptured within a minute, which was followed by a rapid expansion of their chromosomal DNA (Fig. 2a, Supplementary Video 3). Within minutes the DNA expansion reached a stable size (Fig. 2b, c), occupying a typical area of order 50 μm^2^ (or a 3D volume of approximately 80 μm^3^). Lysis method (ii) was performed by flowing a buffer with a low osmolarity (relative to the cell growth medium) through the filling channel of the microfluidic device with trapped spheroplasts. This resulted in a more irregular lysis of the spheroplasts, with some cells lysing but their chromosomal DNA only minimally expanding while others not lysing at all (Supplementary Fig. 4). Therefore, in all the following experiments, we used the surfactant-based lysis. However, as residual IGEPAL can potentially interfere with downstream protein-binding experiments, we explored what minimal concentration could be used to still yield robust lysis. We were able to lyse cells with only 0.2% IGEPAL and adopted that as a working concentration.

**Fig. 2.**
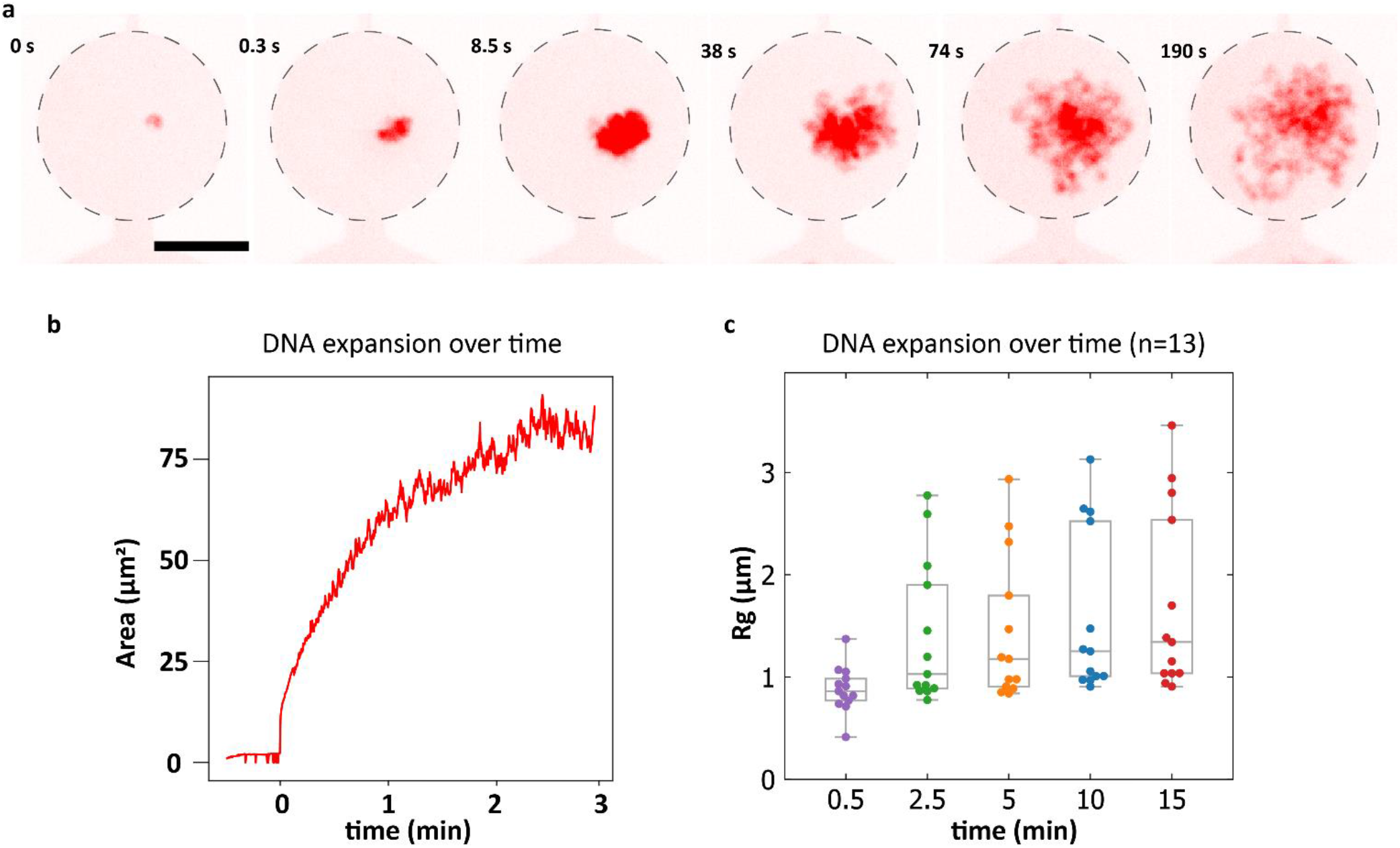
Trapping and lysis of quasi-2D confined *B. subtilis* spheroplasts. **a**. Sequence of images showing the lysis event of a single *B. subtilis* cell and the gradual expansion of the Sytox Orange labeled genomic DNA in the quasi-2D confined environment. Scale bar is 10 μm. **b**. Detected area of the chromosomal DNA from **a** over time. **c**. Calculated radius of gyration of n=13 *B. subtilis* nucleoids over time.

### Upon lysis, proteins dissociate from the megabasepair DNA

As the extracted bacterial genomic DNA is intended to be the starting material for studying the binding of chromosome-organizing proteins to bare DNA, we aimed to remove the original cellular proteins from the nucleoids. Upon lysis, most of these in fact spontaneously unbound from the nucleoid and diffused away. To measure how many proteins remained bound to DNA, we first used an amine-reactive fluorescent dye (Alexa647-NHS) to nonspecifically label the cellular proteins in *B. subtilis*. The succinimidyl ester group on this molecule reacts with primary amines (N-terminus and lysine residues), making all proteins viable targets for labeling. Although we expected this dye to react with cellular proteins only after the cells had been lysed, we did, interestingly, find that the dye was able to already permeate the membrane of the spheroplasts and thus enter the cytoplasm of the spheroplasts and label proteins therein (Fig. 3A). As the cells were lysed, the Alexa647 signal faded away from the DNA within seconds, indicating that the bulk of the *B. subtilis* proteins dissociated from the DNA very rapidly. Since the relatively low signal-to-background ratio of around 5 (Supplementary Fig 5) limited the sensitivity of this assay, we decided to further investigate the degree of protein removal with mass spectrometry (Supplementary Fig. 7). Mass spectrometry samples were prepared in similar conditions in dialysis plugs (Materials and Methods) to mimic what happens in the microfluidic device. The mass spectrometry data indicated that a majority of total protein dissociated from both *E. coli* and *B. subtilis* genomic DNA. In particular, the amount of DNA-binding protein was reduced by at least 10-fold upon treatment (Supplementary Table 1) for *B. subtilis*.

**Fig. 3.**
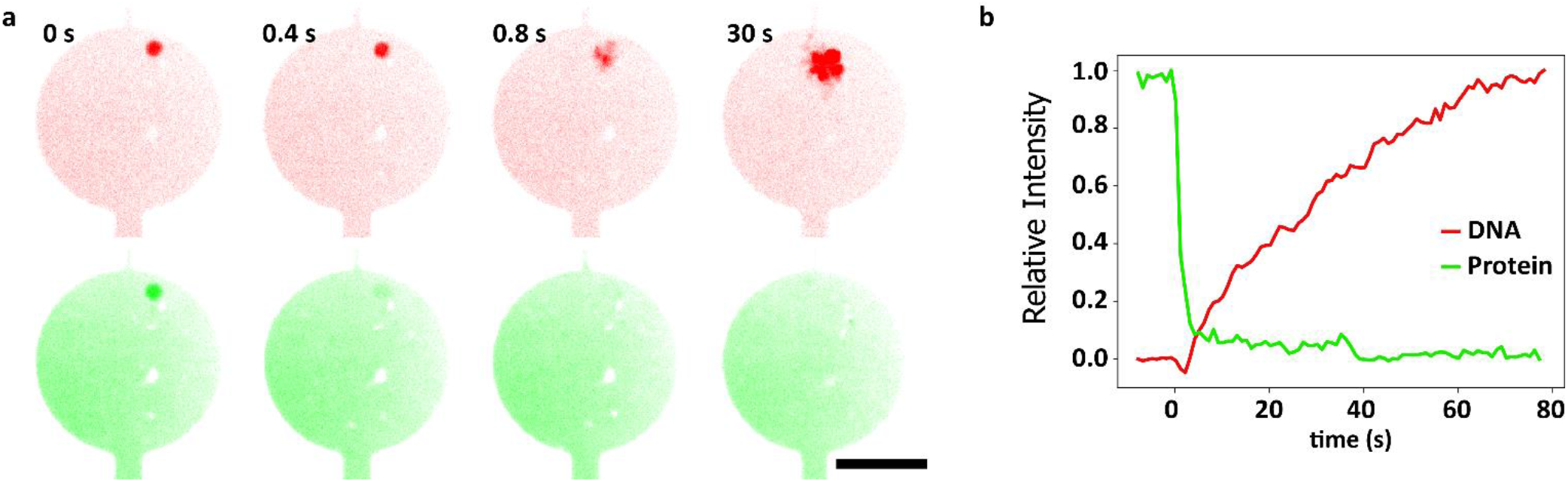
Analysis of the extracted bacterial genomic DNA in microfluidic chambers. **a**. Sequence of confocal fluorescence micrographs showing the lysis event of a single *B. subtilis* spheroplast. DNA (red channel) was labeled using Sytox Orange, while an amine reactive fluorescent label Alexa647-NHS (green channel) was used to track the movement of intracellular proteins during the lysis event. Scale bar is 10 μm. **b**. Normalized fluorescence intensities of the of the *B. subtilis* spheroplast from **a**. Green and red traces correspond to the Alexa647-NHS and Sytox Orange intensities respectively.

### DNA-binding proteins and crowders can condense DNA

To demonstrate the capability of our microfluidic platform to introduce DNA-organizing elements to the trapped megabasepair-long and deproteinated DNA, we probed for the effects of a generic molecular crowding agent (PEG) and of a DNA-binding protein Fis to visualize DNA condensation in real time (Fig. 4a). Introduction of 10% PEG solution into the filling channel of the microfluidic device resulted in the significant compaction of the chromosomal DNA within the trapping chambers (Fig. 4b). We observed that most densely compacted DNA was more prone to adsorption to the walls of the microfluidic chambers (Supplementary Fig. 6). When instead 3 μM of fluorescently labeled Fis protein was flowed into the microfluidic device, we observed its binding to the trapped chromosomal DNA (Fig. 4c). However, in this case, no significant change to the shape or size of the DNA was observed. These results are proof-of-principle illustrations of how our microfluidic platform allows for a diffusion-based addition of DNA-organizing elements to bacterial chromosomal DNA without perturbing the fragile megabase-scale DNA in the process.

**Fig. 4.**
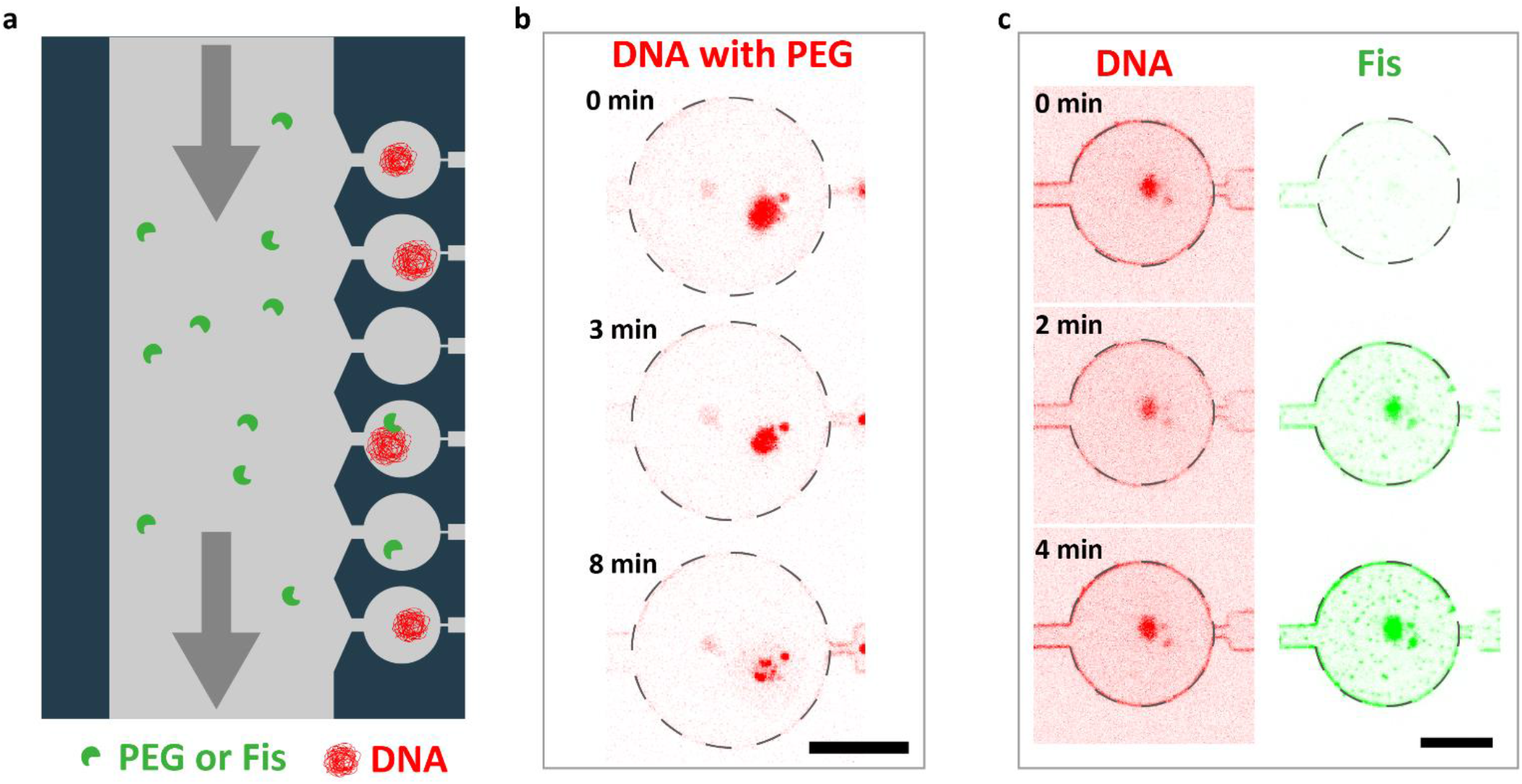
Manipulating the extracted chromosome using DNA-organizing elements. **a**. Experimental setup. DNA-interacting elements are flowed through the filling channel from where they can diffuse into the trapping chambers and interact with the trapped DNA molecules. **b**. A sequence of confocal fluorescence micrographs of a *B. subtilis* nucleoid being exposed to a 10% PEG solution. Scale bar is 10 μm. **c**. A sequence of confocal fluorescence micrographs of a *B. subtilis* nucleoid being exposed to a 3 μM Fis solution. Scale bar is 10 μm.

## Conclusions

We presented a microfluidic platform for the *in vitro* study of genome-sized DNA where DNA-organizing elements can be added without perturbing the trapped genomes. Studying such genome-sized DNA molecules with a “genome-in-a-box” approach^15^ aims to fill the gap between live-cell and single-molecule experiments. A two-layer PDMS chip with integrated valves and cell-trapping chambers was used to trap and subsequently lyse *B. subtilis* spheroplasts, whereupon most of the DNA-binding proteins detached from the nucleoid. The approach allows for the extracted chromosomal DNA to be continuously observed from the moment of cell lysis.

Our work builds on previous studies of isolated *E. coli* nucleoids in bulk solution^24,25^ and cell-sized microchannels.^22^ A key limitation of the bulk methods is that it is very difficult to continuously track the behavior of individual DNA molecules, especially when new reagents are being introduced to the solution which exposes the DNA to mechanical disruption and concentration gradients. The main advantage of our approach compared to microfluidic devices with cell-sized microchannels is that the precise flow control provided by the integrated valves and the ability to direct the fluid flow through the trapping chambers allows for seamless cell loading and introduction of reagents to the trapping chambers. The use of the quasi-2D geometry in 1.6 micrometer high chambers makes it possible to image the isolated chromosomal DNA in a single plane, and resolve its finer structure and dynamics. Most importantly, the approach allows to locally trap a megabasepair-long DNA molecule and subsequently administer new components by diffusion, i.e. not by a flow which disrupts the DNA. Next to the great potential of the methodology, it also has some limitations, for example, some residual undesired surface interactions of the chromosomal DNA at very high densities, and the fact that custom-made microfluidic devices are single-use which leads to a relatively low overall experimental throughput.

Summing up, we developed a cell lysis method using a small amount of surfactant. This led to a rapid expansion of the chromosomal DNA and dissociation of cellular proteins from the DNA. We used mass spectrometry to verify that the DNA is mostly protein-free after this treatment. Proof-of-principle experiments using a crowding agent and DNA-binding protein Fis demonstrated the feasibility of the microfluidic “genome-in-a-box” approach. We envision to use the new microfluidic platform for further bottom-up studies of genome organization. Examples will include the effects of loop-extruding proteins on a genome-sized DNA, behavior of the nucleoid under spatial confinement, and *in-vitro* transcription-translation from genomic DNA.

## Materials and methods

### Microfluidic device fabrication

The PDMS/glass microfluidic devices were fabricated using 2-layer soft-lithography techniques.^23^ The bottom (flow) layer master mold was fabricated using a combination of electron-beam (e-beam), photo-lithography, and DRIE etching. Etching mask for features with 1.6 um height was generated by spin-coating NEB-22 e-beam resist at 1000 rpm for 60 s on a 4” silicon wafer, followed by a 120 s bake at 110 °C. The patterns were then exposed using EBPG-5200 (Raith Nanofabrication), followed by a 120 s bake at 105 °C and developed for 60 s in MF322. The patterns were then DRIE etched into the silicon wafer on Oxford Estrellas using Bosch process at 5 °C in 14 steps. Next the 10 um features with rounded profiles were fabricated by spin-coating AZ10XT positive photoresist at 2000 rpm for 60 s, followed by a 180 s bake at 110 °C. The patterns were then exposed using a Heidelberg uMLA direct writer with a dose of 500 mJ/cm2 and developed for 6 min in AZ400K (diluted 1:3 in demi-water). Rounded profile was then obtained by placing the wafer on a 25 °C hotplate and then ramping the temperature to 120 °C in approximately 5 min, after which the wafer was allowed to cool down by switching off the hotplate. This allows the resist to reflow without introducing cracks or bubbles which often appear when placing a wafer with solidified AZ10XT directly on a 120 °C hotplate.

The top (control) layer master mold was fabricated using photo-lithography and DRIE etching. ARN4400.05 photoresist was spin-coated at 4000 rpm for 60 s on a 4” silicon wafer, followed by a 120 s bake at 90 °C. The patterns were then exposed using a Heidelberg uMLA direct writer with a dose of 60 mJ/cm2, followed by a 5 min bake at 100 °C and developed for 75 s in MF321. The patterns were then DRIE etched 20 μm into the silicon wafer on Oxford Estrellas using Bosch process at 5 °C in 150 steps.

The final devices consisted of bottom (flow) and top (control) layers that were bonded to a glass coverslip. We fabricated the layers with 2-layer soft-lithography techniques using ratios 18:1 and 6:1 of PDMS base to curing agent (Sylgard 184 Silicone Elastomer Kit, Dow Corning GmbH) for the bottom and top layers respectively. PDMS was desiccated before casting over the molds, and the desiccation was repeated for the top layer after casting. The bottom layer was spin-coated at [3’000 rpm for 60 s]. The two layers were baked at 90 °C for around 10 min until the top layer PDMS had hardened while the thin bottom layer PDMS was still slightly sticky to the touch. PDMS slabs were then cut out from the top layer castings and manually aligned and placed on top of the bottom layer. The two PDMS layers were gently pushed together but no weights were used as this often resulted in collapsing the 1.6 μm flow layer channels. Next, the two layers were thermally bonded by baking at 90 °C for 2 to 3 hours. The bonded PDMS devices were gently peeled off the bottom layer wafer, and inlet and outlet holes were manually punched with 0.5 mm diameter biopsy punch. Finally, the PDMS blocks were bonded onto the glass coverslips (#631-0147, 24×50 mm No.1.5, VWR (Avantor) International BV) using oxygen plasma (#119221 Atto, Diener electronic GmbH + Co. KG) at 40 W for 20 s.

### Bacterial cell culture

*E. coli* bacterial cells (BN2179, HupA-mYPet frt, Ori1::lacOx240 frt, ter3::tetOx240 gmR, ΔgalK::tetR-mCerulean frt, ΔleuB::lacI-mCherry frt, DnaC::mdoB::kanR frt)^26^ were incubated from glycerol stock in LB media supplemented with 50 µg/mL Kanamycin antibiotic (K1876, Sigma-Aldrich) in a shaking incubator at 30°C and 300 rpm overnight. The cells were then resuspended in the morning to OD=0.05 and allowed to grow for until reaching OD of 0.1 (approx. 1 hour). The cells were then grown for another hour at 41°C shaking at 900 rpm in order to arrest replication initiation. Next, appropriate volume of cell culture was spun down at 10000 g for 2.5 min, in order to obtain a pellet at OD_eq_ = 1 (approx. 8 × 10^8^ cells). The pellet was resuspended in 475 µL cold (4°C) sucrose buffer (0.58 M sucrose, 10 mM Sodium Phosphate pH 7.2, 10 mM NaCl, 100 mM NaCl). 25 µL lysozyme (L6876 Sigma-Aldrich, 1 mg/mL in ultrapure water) was immediately added and gently mixed into the cell/sucrose buffer suspension, followed by 30+ min incubation at room temperature to create spheroplasts.

*B. subtilis* bacterial cells (BSG4623, smc::-mGFP1mut1 ftsY::ermB, hbsU-mTorquais::CAT, ParB-mScarlet::kan, amyE::Phyperspank-opt.rbs-sirA (spec), trpC2)^27^ were incubated from glycerol stock in SMM+MSM medium (300 mM Na2-Succinate, supplemented with 0.1% Glutamic acid and 2ug-mL Tryptophan) in a shaking incubator at 30°C and 300 rpm overnight. The cells were resuspended in a fresh media in the morning (12.5x dilution of the overnight culture) and allowed to grow for 3 hours. Subsequently, 2 mM IPTG was added to the culture to arrest replication, while continuing shaking at 30°C and 300 rpm. Finally, to create spheroplasts, lysozyme was added to the culture to final concentration of 500 ug/mL for at least 40 minutes. Spheroplasts created in either of two ways were then directly used for on-chip experiments.

### Expression, purification, and labelling of Fis

Full length *Escherichia coli* Fis with an N-terminal His_8_ tag followed by a HRV-3C protease site, and appended with a C-terminal cysteine residue, was expressed from (pET28a-derived) plasmid pED72 in *Escherichia coli* ER2566 cells (New England Biolabs, *fhuA2 lacZ::T7 gene1 [lon] ompT gal sulA11 R(mcr73::miniTn10--Tet*^*S*^*)2 [dcm] R(zgb-210::Tn10--Tet*^*S*^*) endA1 Δ(mcrCmrr)114::IS10)*. Cells were grown at 37 °C in baffled flasks on LB supplemented with 50 µg/ml kanamycin, expression was induced at an OD_600_ of 0.6 with 0.2 mM IPTG, and cells were harvested after overnight expression at 18 °C (8 min 4500 rpm, JLA8.1000 rotor). After washing the cells in PBS they were resuspended in buffer A (50 mM TrisHCl pH 7.5 (@RT), 750 mM NaCl, 1 mM EDTA, 0.05 mM TCEP, 10% (w/v) glycerol) and lysed using a French Press (Constant Systems) at 20 kpsi, 4°C. Following the addition of 0.35% polyethyleneimine, unbroken cells, DNA and protein aggregates were pelleted in a Ti45 rotor (30 min, 40.000 rpm, 4 °C), and Fis was precipitated from the supernatant by the addition of 476 g/l ammonium sulfate. Following centrifugation (JA-17 rotor, 10 minutes, 8500 rpm, 4 °C) and resuspension in buffer A, the sample was applied to 2 ml Talon Superflow resin (Clontech) pre-equilibrated with buffer A, and incubated for one hour while rotating at 4 °C. Subsequently, the resin was washed with buffer A supplemented with 20 mM imidazole and finally Fis was eluted in 15 ml of buffer A supplemented with 1 mM β-mercaptoethanol and homemade 3C protease. Proteins were concentrated using a Vivaspin centrifugal concentrator (10 kDa cut-off) and further purified by size exclusion chromatography (SEC) on a Superdex 200 Increase 10/300 column pre-equilibrated with buffer A, eluting at ∼16.5 ml. For preparation of fluorescently labelled Fis, 0.5 ml of concentrated protein was incubated 0.1 mM Alexa Fluor™ 647 C2 Maleimide (Invitrogen) for 30 minutes at room temperature, prior to size exclusion chromatography. Purified protein was snap-frozen and stored at -80 °C until use.

### Operation of the microfluidic nucleoid trapping and analysis device

The microfluidic device was mounted on the stage of a spinning disk confocal microscope (Andor …). The operation of the devices requires precisely controlling pressure on the input lines, as well as supplying steady pressure on the valve lines. The control/valve channels of the device were filled with MilliQ water and actuated using a pneumatic valve array (FESTO), which was in turn actuated using an array of manual switches connected to a benchtop power-supply. The input pressure to the pneumatic valve array was 2 bar. The pressure to the reagent input channels of the microfluidic device was controlled using an adjustable pressure regulator (Fluigent). In a typical experiment, buffer solution (20 mM Tris-HCl ph 7.5, 50 mM NaCl, 1 mg/ml BSA) was connected to inlet 1 of the microfluidic device with a pressure of 300 mbar in order to wet all the flow channels and remove any air bubbles. Next the trapping chamber area of the device was filled with a buffer containing DNA intercalating dye (Sytox Orange, 400 nM) and incubated for 15 minutes.

As a next step, the bacterial spheroplasts should be trapped in the microfluidic chambers. To do so, they were injected into the device from inlet port 1 or 2, typically an input pressure of 1-5 mbar was used. Initially the spheroplast were added to the large filling channel by opening valves 1 (or 2), 5 and 7. After a sufficient number of spheroplasts were present in the filling channel, valve 7 was closed and valves 8 and 9 were opened to enable flow through the exhaust channels and thereby allow the spheroplasts to enter the trapping chambers. When a desired amount of spheroplasts had entered the chambers, valves 8 and 9 were closed and at this point the cells were ready for lysis. To lyse the spheroplasts, lysis buffer (Tris-HCl pH 7.5 40 mM, Potassium Glutamate 50 mM, BSA, 0.2 mg/mL, MgCl2 2.5 mM, Glucose 5%, Sytox Orange 500 nM, with addition of IGEPAL-CA-630 0.2% to aid lysis) was connected to inlet port 3 and was injected into the filling chamber by opening valves 3, 6 and 7 and using an input pressure of 1-2 mbar. Lysis of the individual spheroplasts could then be observed, this proceeded in a sequential manner starting from the upper trapping chambers. Stopping the flow of the lysis buffer would also stop the lysis events from happening in the downstream chambers and this allowed us to analyze the expansion of several nucleoids sequentially with a high frame rate within the same experiment. After all the spheroplasts had been lysed, valves 6 and 7 were closed and a desired reagent (PEG or Fis solution in this case) was connected to inlet 4. Valves 6 and 7 were then reopened and the reagent solution was allowed to flow into the filling channel and to diffuse into the trapping chambers and interact with the trapped DNA.

### Image acquisition and analysis

To image isolated nucleoids in microfluidic traps, we used an Andor Spinning Disk Confocal microscope equipped with 100x magnification oil immersion objective. Isolated DNA was labelled by the intercalating dye Sytox Orange (S11368, Thermo Fischer Scientific, MA, USA) at concentration of 500 nM. At this concentration, the dye is known to reduce the persistence length of DNA to 37 nm. The dye was excited with 561 nm laser line (20% power, 250x gain, 10 ms exposure) with 617/73 nm filter on the emission. The acquisition computer was running Andor iQ 3.6 software. Multiple z-planes per each object, with separation of 1 µm between subsequent planes were acquired. For extended observations, we defined xy-positions and imaged them repeatedly over time, usually once every 30 or 60 seconds.

The analysis of nucleoid images within microfluidic traps was conducted using a custom Python code pipeline. We began by selecting circular regions of interest from in-focus plane images, encompassing the area inside the traps. These image sections were then thresholded to eliminate background noise and isolate the pixels containing fluorescent signal associated with nucleoids. The resulting set of pixels, each characterized by [position, intensity] values, was used to compute the radius of gyration for each nucleoid. This same pixel set also provided a measure of the total thresholded area occupied by the nucleoid.

### Sample preparation for mass spectrometry

Dialysis plug were chosen for sample preparation as they allowed to continuously exchange solutions in which nucleoid were suspended, similar to what happens in the microfluid device. This approach also allowed for removal of IGEPAL, which is otherwise incompatible with LC/MS, even at small concentrations [ref]. Spheroplasts were prepared from overnight cultures as described in the section ‘Bacterial cell culture’. Lysis buffer contained final concentration of 0.2% IGEPAL and 50 mM Tris-HCl (pH 8). All spheroplast samples were lysed by adding 100 uL of spheroplast suspension to 900 uL of lysis buffer in dialysis plugs. Control and treatment samples were prepared in 3.5-5 kDa (G235029, Repligen Corporation, CA USA), and 300 kDa cut-off plugs (G235036, Repligen Corporation, CA USA) respectively following manufacturer’s protocol. Each sample condition was prepared and measured in triplicates.

100 mM ammonium bicarbonate buffer (ABC) was prepared by dissolving ammonium bicarbonate powder (A6141, Sigma-Aldrich) in LC-MS grade quality water. 10 mM DTT (43815, Sigma-Aldrich) and iodoacetamide (IAA) (I1149, Sigma-Aldrich) solutions were made fresh by dissolving stock powders in 100 mM ABC. Next, 50 µL of 100 mM ABC buffer was added to 200 uL of each sample to adjust pH, immediately followed by addition of 60 µL of 10 mM DTT and 1 hour incubation at 37°C and 300 rpm in dark. Next, 60 µL of 20 mM IAA was added and samples were incubated in dark at room temperature for 30 min. Finally, 20 µL of 0.1 mg/mL trypsin (V5111, Promega) was added and samples were incubated for 16-20 hours at 37°C and 300 rpm. On the following day, samples were purified by solid phase extraction (SPE). SPE cartridges (Oasis HLB 96-well μElution plate, Waters, Milford, USA) were washed with 750 µL of 100% methanol and equilibrated with 2×500 µL LC-MS grade H_2_O. Next, 200 µL of each sample was loaded to separate SPE cartridge wells and wells were washed sequentially with 700 µL 0.1% formic acid, 500 µL of 200 mM ABC buffer and 700 µL of 5% methanol. Samples were then eluted with 200 µL 2% formic acid in 80% methanol and 200 µL 80% 10 mM ABC in methanol. Finally, each sample was collected to separate low-binding 1.5 µL tubes and speedvac dried for 2-3 hours at 45°C. Samples were stored frozen at -20°C until further analysis. Desalted peptides were reconstituted in 15 µL of 3% acetonitrile/0.01% formic acid prior to mass spectrometric analysis. Per sample, 2 µL of protein digest was analyzed using a one-dimensional shotgun proteomics approach^28,29^. Briefly, samples were analyzed using a nano-liquid-chromatography system consisting of an EASY nano LC 1200, equipped with an Acclaim PepMap RSLC RP C18 separation column (50 μm x 150 mm, 2 μm, Cat. No. 164568), and a QE plus Orbitrap mass spectrometer (Thermo Fisher Scientific, Germany). The flow rate was maintained at 350 nL/min over a linear gradient from 5% to 35% solvent B over 90 min, then from 35% to 65% over 30 min, followed by back equilibration to starting conditions. Data were acquired from 0 to 130 min. Solvent A was H2O containing 0.1% FA and 3% ACN, and solvent B consisted of 80% ACN in H2O and 0.1% FA. The Orbitrap was operated in data-dependent acquisition (DDA) mode acquiring peptide signals from 385–1250 m/z at 70,000 resolution in full MS mode with a maximum ion injection time (IT) of 75 ms and an automatic gain control (AGC) target of 3E6. The top 10 precursors were selected for MS/MS analysis and subjected to fragmentation using higher-energy collisional dissociation (HCD). MS/MS scans were acquired at 17,500 resolution with AGC target of 2E5 and IT of 100 ms, 2.5 m/z isolation width and normalized collision energy (NCE) of 28.

Mass spectrometric raw data were analyzed against the proteome database from *Escherichia coli* K12 (UP000000625, Tax ID: 83333, April 2024) or *Bacillus subtilis* strain 168 (UP000001570, Tax ID: 224308, April 2024, downloaded from https://www.uniprot.org/)^30^ using PEAKS Studio X (Bioinformatics Solutions Inc., Waterloo, Canada)^31^ allowing for 20 ppm parent ion and 0.02 m/z fragment ion mass error, 3 missed cleavages, carbamidomethylation as fixed and methionine oxidation, N/Q deamidation and N-terminal Acetylation as variable modifications. Peptide spectrum matches were filtered for 1% false discovery rates (FDR) and identifications with ≥ 1 unique peptide matches. The protein area was determined from the averaged top-3 peptide areas. Protein areas between conditions were compared by label free quantification using PEAKSQ, allowing a retention time shift tolerance of 5.0 minutes, a mass error tolerance of 10.0 ppm, and considering protein identifications filtered for 1% FDR. Peptide ID counts and min confident samples was set to 0 and significance method was set to ANOVA. Otherwise software default parameters were used. Data inspection revealed that one *B. subtilis* treatment sample was indistinguishable from the control, and highly dissimilar to other two treatment samples. This pointed to an experimental error and this sample was left out from further analysis.

Relative protein abundancies were defined as the ratio of the ‘treatment’ over the ‘control’ conditions for the top-3 peptide areas, where the areas were weighted by each protein’s molecular mass. For purposes of plotting, where no protein was identified on treatment condition, the fold change was set to 10^−3^, and where no protein was measured on control condition, the fold change was set to the highest one in the dataset. Similarly fold change was limited between 2^7^ and 2^−7^ and plotted as log2(FC) (e.g. log2(2^7^) = 7), and the maximum significance was capped at 10^−20^ (i.e. -log10(10^−20^) = 20) for visualization purposes. To calculate the ratio between conditions presented in Supplementary Table S2, the top-3 peptide areas were summed up per each sample, and the values aggregated per each condition. Standard error of the mean from each condition was propagated to the error on the ratio by propagation of uncertainty.

In this study, we conducted a label-free quantification to compare the control with the purified sample. It is important to note that in such an experiment the remaining proteins in the purified sample are expected to appear more abundant than when they are part of a complex mixture. As a result, the apparent abundance of these proteins may seem higher in the purified sample compared to the control. The relative abundance of proteins after purification, as reported in Supplementary Table S2, should therefore be considered an upper bound estimate, and the actual quantities are likely significantly lower.

## Acknowledgements

The authors would like to acknowledge Miloš Tišma and Jaco van der Torre for their help and advice on culturing the *B. Subtilis* BSG4623. Dita Heikens is acknowledged for her assistance with the proteomics experiments. We acknowledge funding support from the ERC Advanced Grant 883684.

