## Supplemental Information (Figures S1-S7, Tables S1-S3) for "A microfluidic platform for extraction and analysis of bacterial genomic DNA"

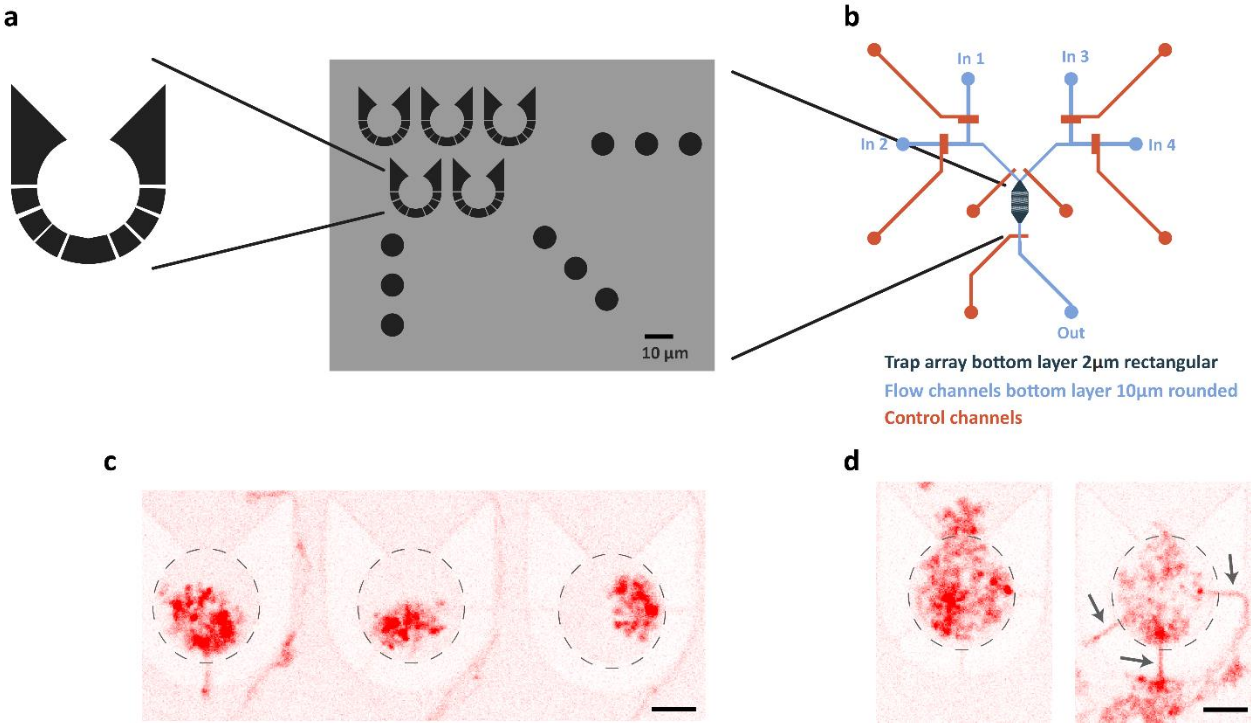


**Supplementary Figure 1.** Initial design of the microfluidic trapping device for bacterial chromosome extraction. **a**. Configuration of a flow cell with a 2D grid of PDMS traps. **b**. Schematic of the complete microfluidic chip. The left inputs (in1 and in2) were used for loading the cells while the right inputs (in3 and in4) were used for loading the lysis buffer. **c**. Confocal fluorescence micrograph of extracted genomic DNA from 3 *B. subtilis* cells labeled with Sytox Orange. Scale bar is 5 µm. **d**. An example of DNA leakage through the narrow slits of the PDMS trap. Scale bar is 5 µm.

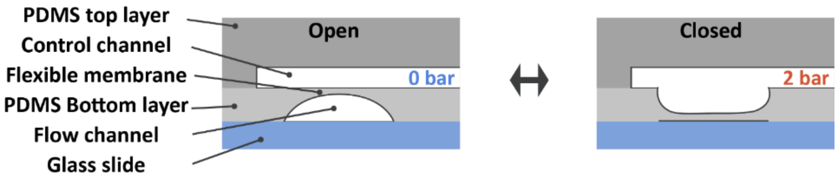


**Supplementary Figure 2.** Design and fabrication of pneumatically actuated microfluidic valves. Push-down configuration was used for all the valves. The rounded profile in the mold for valves 1, 2, 3, and 4 was realized by reflowing AZ10XT photoresist at 120C.

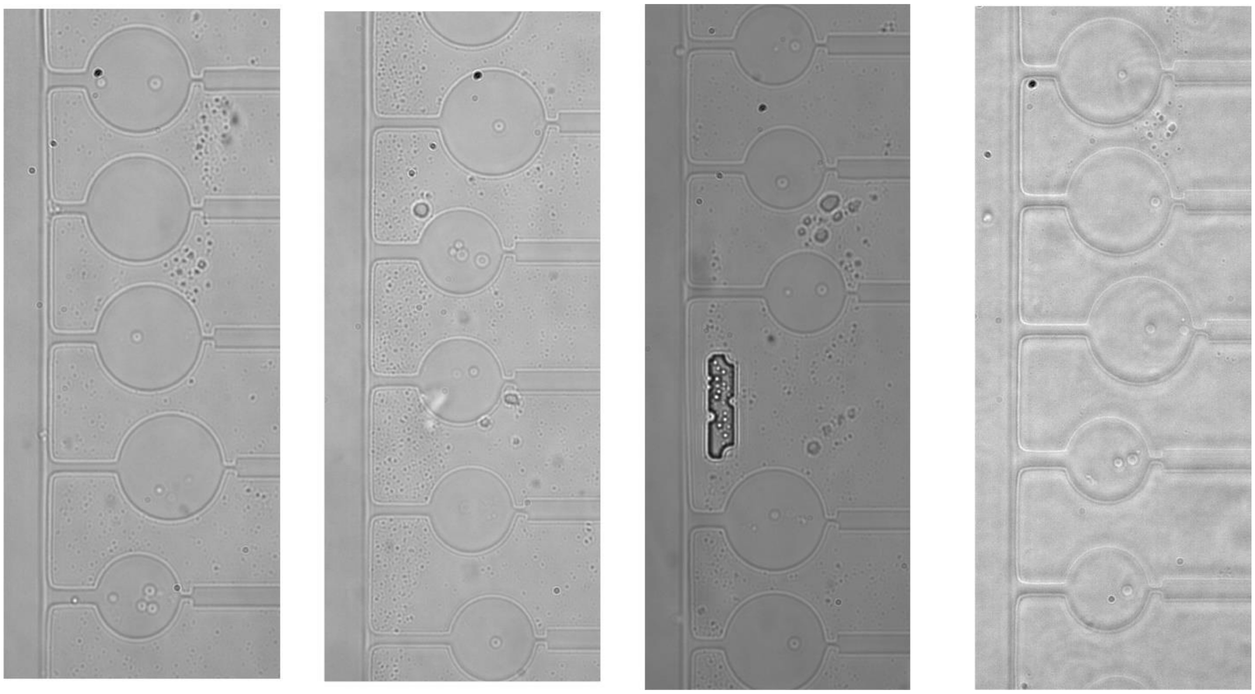

**Supplementary Figure 3.** Widefield micrographs of *B. subtilis* spheroplasts in the microfluidic trapping chambers. Approximately 40% of the traps contain a single spheroplast.

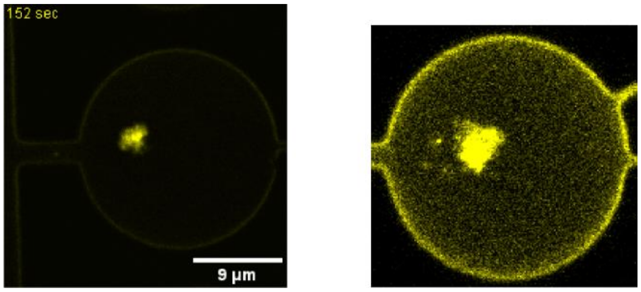


**Supplementary Figure 4.** Lysis results using osmotic shock. Two examples of Sytox-Orange-labeled genomic DNA that was extracted using an osmotic shock, resulting in a more compacted structure.

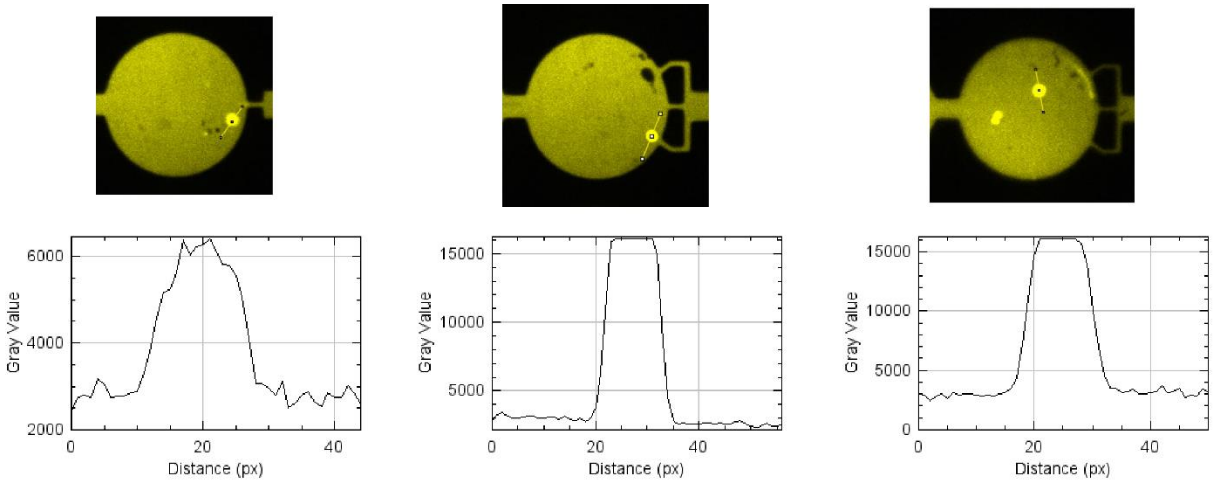


**Supplementary Figure 5.** Signal-to-background ratio of Alexa647-NHS protein quantification measurements. Three examples of Alexa647 fluorescence intensity profiles of spheroplasts before lysis.

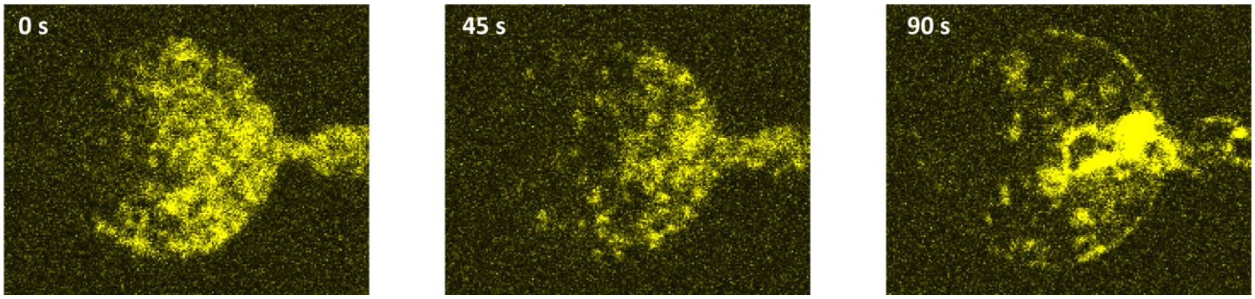


**Supplementary Figure 6.** Adsorption of condensed DNA to chamber walls. An example of DNA condensation and absorption to the bottom of the trapping chamber upon addition of 20% PEG solution.

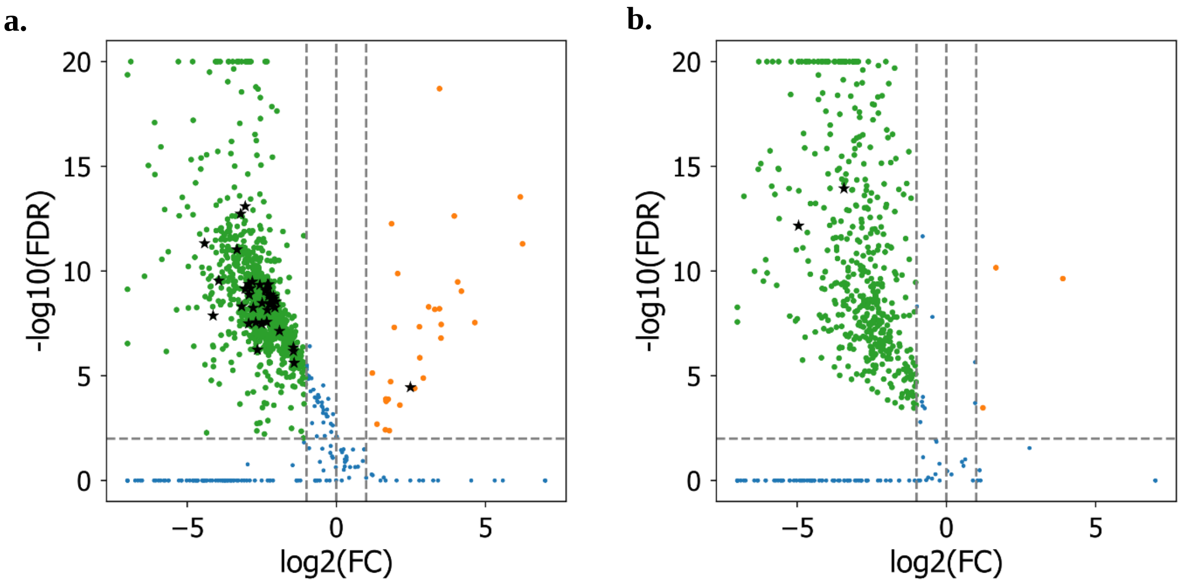


**Supplementary Figure 7:** Protein abundance after removal for **a**. *E. coli* **b**. *B. subtilis*. Vertical lines indicate 2-fold removal and enrichment respectively, horizontal line corresponds to significance threshold of 1%. Points highlighted with stars represent DNA-binding proteins (cf. Table S2 and S3).

| | ***E. coli*** | ***B. subtilis*** |
|---|---|---|
| **Relative abundance [%]** | | |
| DNA-binding | 19.2 ± 6.6 | 9.9 ± 2.6 |
| All | 24.6 ± 8.5 | 17.7 ± 4.4 |
| **Number of proteins reduced fewer than two-fold** | | |
| DNA-binding | 1‡ (of 39) | 0 (of 2) |
| All | 69 (of 1246) | 13 (of 490) |

**Table S1:** Relative protein abundance after treatment for *E. coli* and *B. subtilis,* and the number of proteins that were not reduced more than 2-fold. Proteins were selected on significance threshold of 1% on the fold-change. To obtain relative values, proteins were weighted by their mass. ‡ RpoZ

| Protein Name | Description |
|---|---|
| hbs | DNA-binding protein HU 1 |
| rpoY | DNA-directed RNA polymerase subunit epsilon |

**Table S2:** Proteins labeled as DNA-binding or DNA-processing in the *B. subtilis* sample (filtered on FDR 1%).

| Protein Name | Description |
|---|---|
| dnaE | DNA polymerase III subunit alpha |
| topA | DNA topoisomerase 1 |
| cbpA | Curved DNA-binding protein |
| dps | DNA protection during starvation protein |
| matP | Macrodomain Ter protein |
| gyrA | DNA gyrase subunit A |
| mukE | Chromosome partition protein MukE |
| rpoB | DNA-directed RNA polymerase subunit beta |
| ybaB | Nucleoid-associated protein YbaB |
| hupA | DNA-binding protein HU-alpha |
| parC | DNA topoisomerase 4 subunit A |
| mukF | Chromosome partition protein MukF |
| ybiB | Uncharacterized protein YbiB |
| hupB | DNA-binding protein HU-beta |
| mukB | Chromosome partition protein MukB |
| crp | DNA-binding transcriptional dual regulator CRP |
| rpoC | DNA-directed RNA polymerase subunit beta' |
| ompR | DNA-binding dual transcriptional regulator OmpR |
| ihfB | Integration host factor subunit beta |
| rpoA | DNA-directed RNA polymerase subunit alpha |
| rpoD | RNA polymerase sigma factor RpoD |
| crl | Sigma factor-binding protein Crl |
| dnaN | Beta sliding clamp |
| ihfA | Integration host factor subunit alpha |
| polA | DNA polymerase I |
| fis | DNA-binding protein Fis |
| uvrA | UvrABC system protein A |
| gyrB | DNA gyrase subunit B |
| nadR | Trifunctional NAD biosynthesis/regulator protein NadR |
| rpoS | RNA polymerase sigma factor RpoS |
| yaaA | DNA-binding and peroxide stress resistance protein YaaA |
| uvrD | DNA helicase II |
| yejK | Nucleoid-associated protein YejK |
| oxyR | DNA-binding transcriptional dual regulator OxyR |
| stpA | DNA-binding protein StpA |
| parE | DNA topoisomerase 4 subunit B |
| hns | DNA-binding protein H-NS |
| kdgR | HTH-type transcriptional regulator KdgR |
| rpoZ | DNA-directed RNA polymerase subunit omega |

**Table S3:** Proteins labeled as DNA-binding or DNA-processing in the *E. coli* sample (filtered on FDR 1%).
